# Tubulin acetylation increases cytoskeletal stiffness to regulate mechanotransduction in striated muscle

**DOI:** 10.1101/2020.06.10.144931

**Authors:** Andrew K. Coleman, Humberto C. Joca, Guoli Shi, W. Jonathan Lederer, Christopher W. Ward

**Affiliations:** Center for Biomedical Engineering and Technology, University of Maryland School of Medicine, Baltimore, Maryland; Department of Orthopedics, University of Maryland School of Medicine, Baltimore, Maryland

**Author notes:** Contributed equally to the work (Andrew K. Coleman, Humberto C. Joca). Corresponding Authors: H. C. Joca: Centre for Biomedical Engineering and Technology, University of Maryland School of Medicine, Baltimore, MD, USA.; C. W. Ward: Department of Orthopedics, University of Maryland School of Medicine, Baltimore, MD, USA.

## Abstract

Microtubules tune cytoskeletal stiffness to regulate the mechanics and mechanotransduction of striated muscle. While recent evidence suggests that microtubules enriched in detyrosinated α-tubulin are responsible for these effects in healthy muscle, and for their excess in disease, the possible contribution from several other α-tubulin modifications has not been investigated. Here we used genetic or pharmacologic strategies in isolated cardiomyocytes or skeletal myofibers to increase the level of acetylated α-tubulin without altering the level of detyrosinated α-tubulin. We show that microtubules enriched in acetylated α-tubulin contribute to the cytoskeletal stiffness and viscoelastic resistance, showing slowed rates of contraction and relaxation during unloaded contraction, and increased activation of NADPH Oxidase 2 (Nox2) by mechanotransduction. Together these findings add to growing evidence that microtubules contribute to the mechanobiology of striated muscle in health and disease.

## Introduction

Microtubule binding to actin, intermediate filaments and protein cytolinkers regulates cytoskeletal mechanics and mechanotransduction^1–4^. This occurs in striated muscle where there is evidence that disease-altered microtubules contribute to the pathological increase in cytoskeletal mechanics^3–8^ and mechanotransduction^3,4,8^. This pathologic excess of microtubule dependent mechanotransduction manifested in increased susceptibility to contraction induced injury in skeletal muscle and a predisposition to arrhythmic events in the heart^8^. Here the focus has been on function-altering post-translational modifications (PTMs) of α-tubulin by acetylation and detyrosination^9^.

Microtubules are dynamic polymers of α-β protein dimers where detyrosination (deTyrMT) or acetylation (acetylMT) increases filament longevity and binding^9–12^. In striated muscle both deTyrMT and acetylMT are evident in healthy cardiac and skeletal muscle and both are dramatically elevated in elevated in disease. Recent works find deTyrMTs essential for striated muscle myocyte mechanics and mechanotransduction, independent of acetylMTs, and for their excess in disease^8,13^. Despite evidence in neurons that acetylMT’s are a positive regulator of cytoskeletal mechanics and mechanotransduction^14^, recent work in cardiomyocytes suggests a suppressive effect^15^. In any case, the independent role of α-tubulin acetylation on cellular mechanics and mechanotransduction has yet to be been defined. Here we sought to determine the independent role of acetylMTs on cytoskeletal mechanics and mechanotransduction in striated muscle.

## Results and Discussion

The acetylation of α-tubulin is by α-tubulin acetyltransferase 1 (αTAT1) at K40 on a loop (residues P37 to D47)^16,17^ within the microtubule lumen, and reversed by histone deacetylase 6 (HDAC6) or the NAD-dependent deacetylase Sirtuin 2^18,19^. In enzymatically isolated ventricular cardiomyocytes and *Flexor digitorum brevis* (FDB) skeletal myofibers we show that tubacin, a pharmacologic HDAC 6 inhibitor, increased the level of α-tubulin acetylation without altering the level of detyrosinated α-tubulin nor the expression of α-tubulin protein (Fig. 1 A and B). A further examination of microtubule structure in both cell types revealed that tubacin treatment increased the level of tubulin acetylation without altering microtubule density (Fig. 1 C and D). Using this treatment, we can accurately assess the contribution of acetylated tubulin to cellular viscoelastic resistance and microtubule dependent mechano-signaling pathways independent of changes in detyrosinated tubulin or microtubule density.

**Figure 1.**
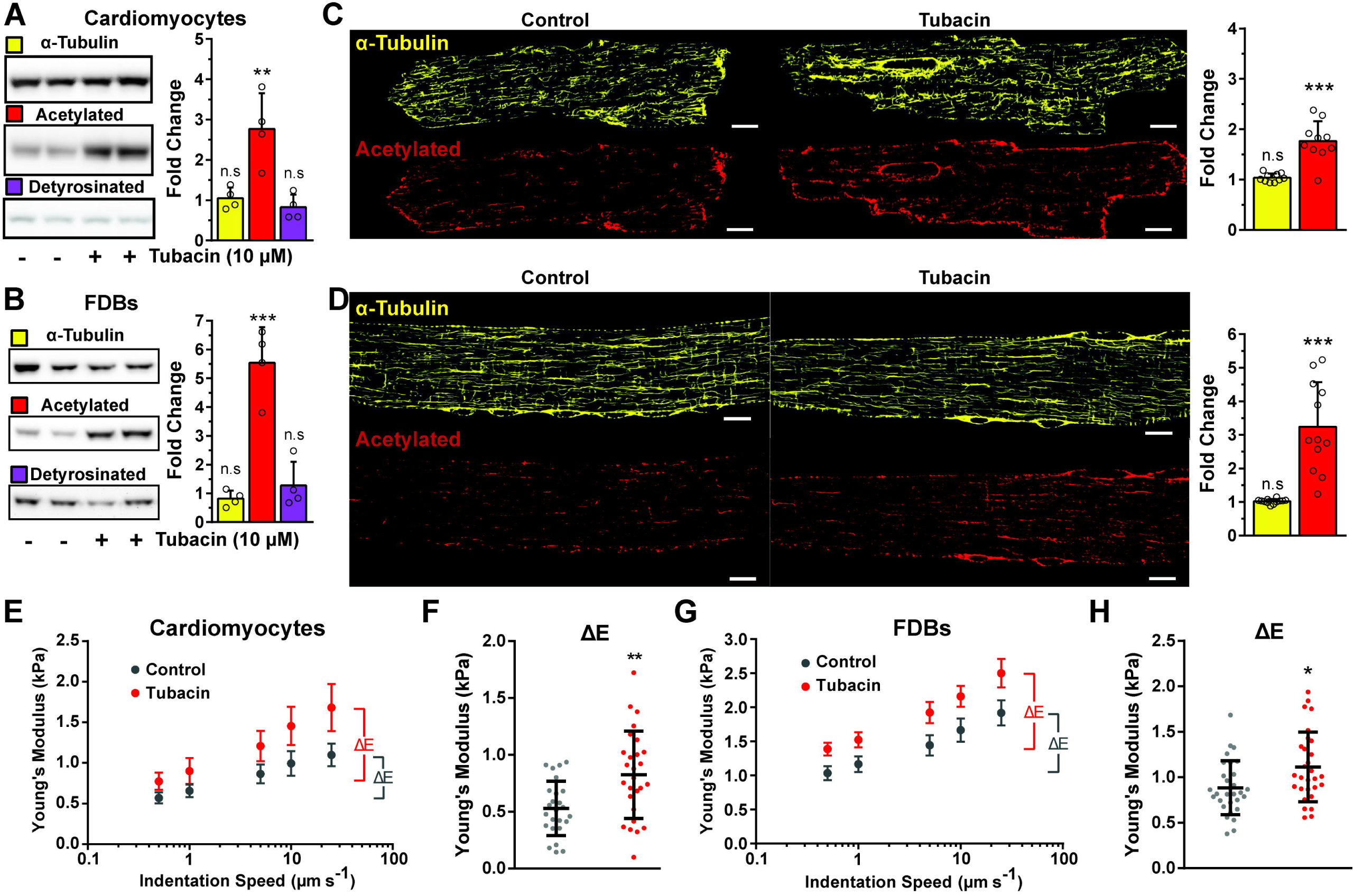
HDAC6 inhibition increases α-tubulin acetylation independent of altering the density of microtubules and increases cell viscoelastic resistance. **A** and **B**) Western blot of isolated cardiomyocytes and FDB fibers treated with HDAC6 inhibitor Tubacin (10 µM, 2h, n = 4). Quantification of α-tubulin, acetylated, and detyrosinated tubulin normalized to ponceau and expressed as fold change to control conditions (vehicle-only). **C** and **D**) Immunofluorescence and quantification of isolated cardiomyocytes and FDB fibers, respectively. Tubulin abundance was determined from a binary quantification normalized to cell area and expressed as fold change to the control. (FDB n = 12, cardiomyocytes, n = 10). Young’s modulus of cardiomyocytes and FDB fibers treated with Tubacin (Red) from nano-indentation at different speeds (**E** and **G** respectively). The estimated viscoelastic resistance from experiments shown in E and G is represented in panels **F** and **H**, respectively. * p<0.05, ** p<0.01, *** p<0.001, scale bars = 10µm.

The contribution of microtubules to the cytoskeletal stiffness and viscoelastic resistance of cardiac and skeletal myocytes is reflected in measures of their transversal stiffness and contractile kinetics^8,20^. While the level of α-tubulin modification by detyrosination has been shown to alter cytoskeletal stiffness and myocyte mechanics, the independent effect of α-tubulin acetylation is unknown. Using measures of nanoindentation mechanics at varying speeds, we show that the tubacin-dependent increase in acetylMTs significantly increases cellular stiffness (Fig. 1E and G) and viscous resistance in both cardiomyocytes and FDB myofibers (Fig. 1F and H). Aligned with these changes, we show a significant reduction in the fractional shortening (Fig. 2 B and F) and rates of both contraction and relaxation (Fig. 2 C and G) during unloaded contractions.

**Figure 2.**
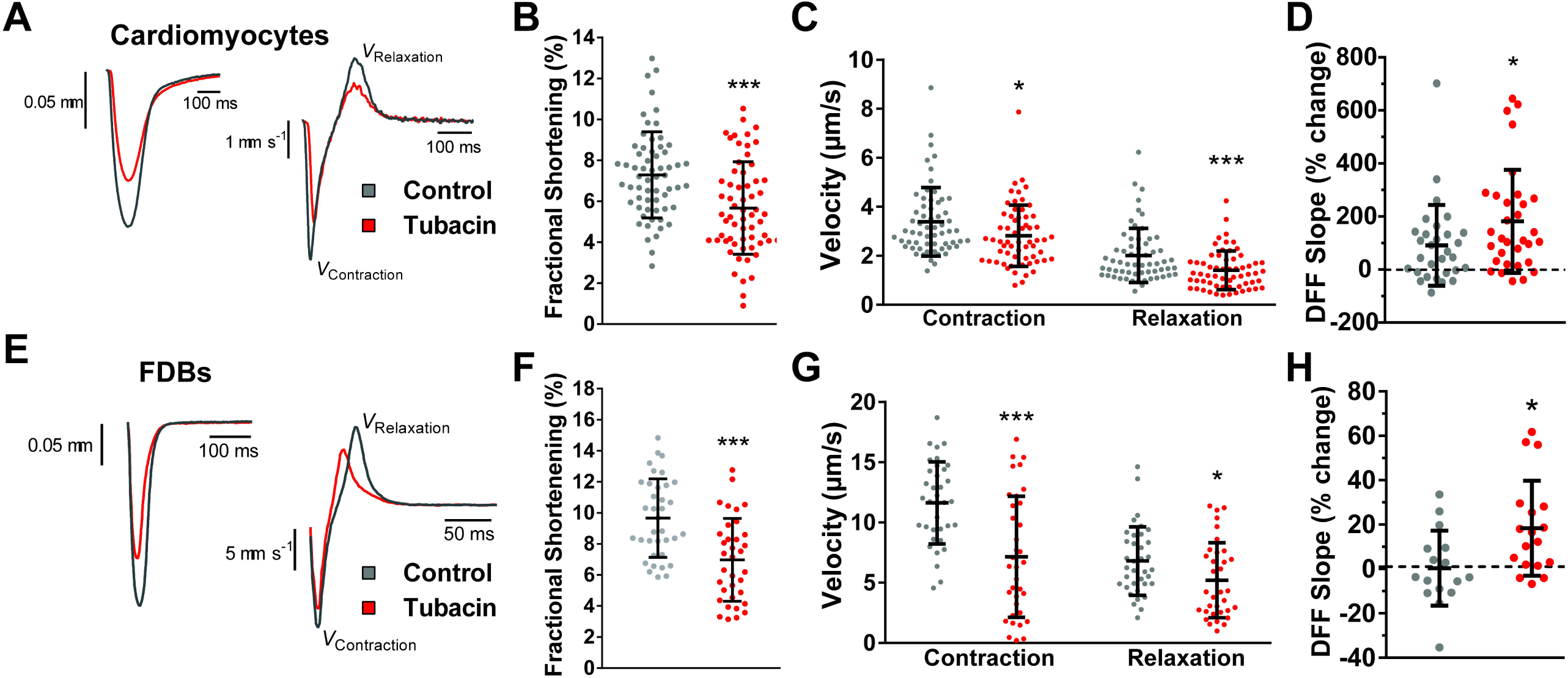
Increased MT acetylation slows contractile kinetics and modulates mechanosensitive ROS production. Representative sarcomere length and velocity during contraction (**A** and **E**) treated with Tubacin (Red) and respective controls (Grey); measured fractional shortening (**B and F**) and contractile kinetics (**C and G**) from cardiomyocytes and FDB fibers, respectively. Panel **D** shows quantification of the change in ROS production in cardiac cells during 2Hz stimulated contractions measured by the change in DFF fluorescence increase and Panel **H** shows quantification of the change in ROS production during 2 Hz cyclic stretch in FDBs fibers. * p<0.05, ** p<0.01, *** p<0.001.

Consistent with the role of microtubules in mechanotransduction^3^, we show a significant increase in mechano-elicited ROS production in single cardiomyocytes undergoing unloaded paced contraction at 2 Hz (Fig. 2 D) and in single FDB’s challenged with 2Hz cyclic passive stretch (Fig. 2 H). These results align with reports that acetylMTs are responsible for the mechanosensitive touch response in mice and Drosophila^14,21^ as well as evidence that acetylated α-tubulin underlies the increased resistance seen with repetitive strain to microtubules *in vitro*^22,23^; a property that would facilitate the transfer of energy for mechanotransduction in striated muscle^4,8,20^.

The abundance of tubulin acetylation is governed by the equilibrium balance of cytosolic deacetylases (including HDAC6) and the acetyltransferase (αTAT1 in mice)^7^ with increased αTAT1 activity mediating the increased α-tubulin acetylation during acute cell stress^24^. As an orthogonal approach to tubacin treatment we used the genetic overexpression of αTAT in mouse FDB muscle in order to induce an increase tubulin acetylation via acetyltransferase mechanism. Following enzymatic isolation of FDB myofibers, we observed an increased level of acetylMTs (Fig. 3A). Measures of passive cell mechanics showed these acetylMTs increased cell stiffness and viscoelastic resistance (Fig. 3B and C) with unloaded shortening confirming the suppressive effect on contractile kinetics (Fig. 3D and E). Those changes in contraction and cellular stiffness fell in line with HDAC6 inhibition approach, which consolidate the impact of this PTM on striated muscle function.

**Figure 3.**
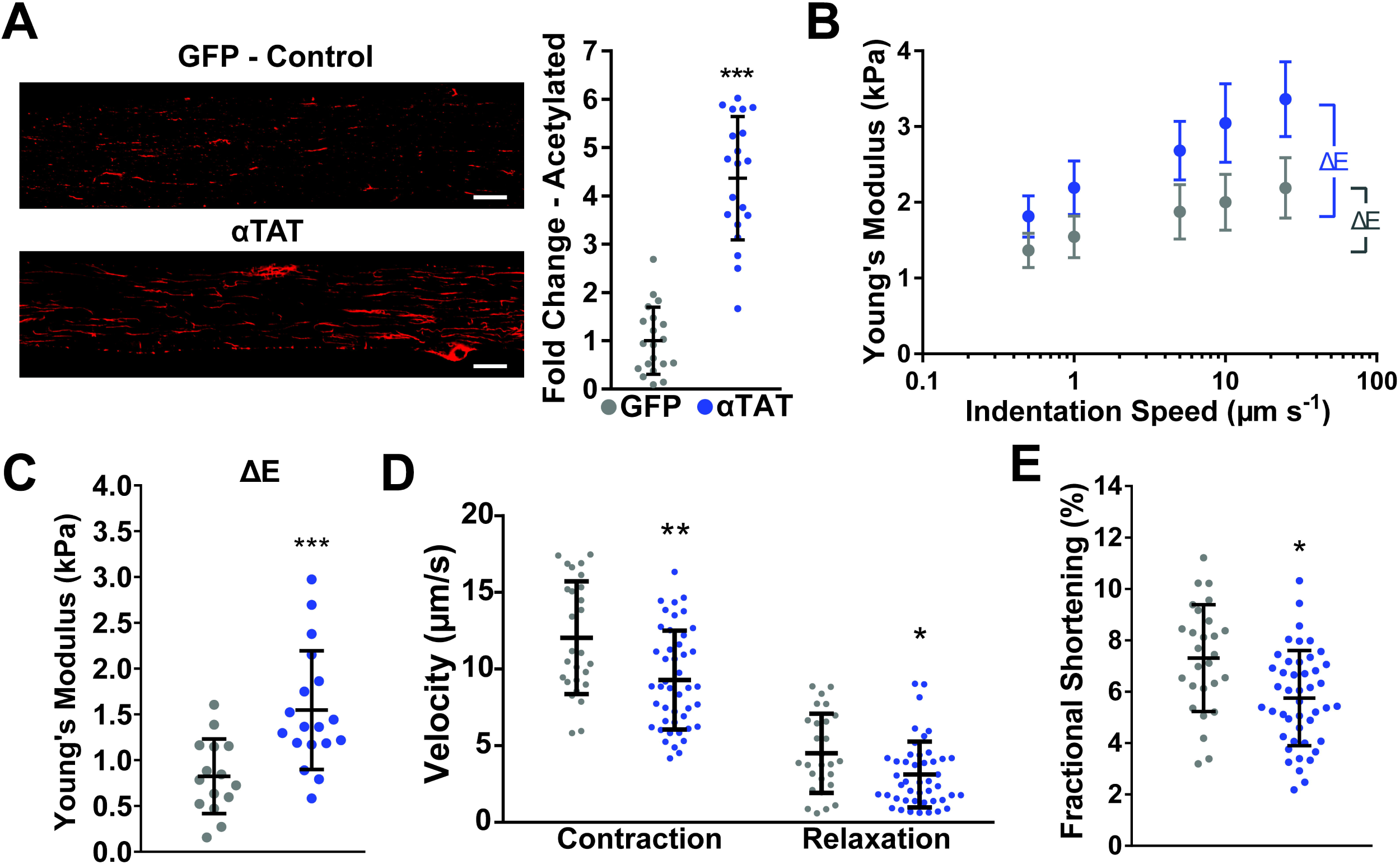
Overexpression of α-tubulin acetyltransferase (αTAT) increases viscoelastic resistance and slows contractile kinetics. **A**) Immunofluorescent staining and quantification of acetylated tubulin in FDBs overexpressing αTAT (fold change to GFP-only controls). Young’s modulus of cardiomyocytes and FDB fibers overexpressing αTAT (Blue) and GFP-control (Grey) from nano-indentation at different speeds (**B**) and their derived viscoelastic resistance (**C**). Peak contraction, relaxation velocities (**D**), and Fractional shortening (**E**) measured from FDB fibers over-expressing αTAT (Blue) and their GFP-only controls (Grey). * p<0.05, ** p<0.01, *** p<0.001, scale bars = 10µm.

Taken together we report that α-tubulin acetylation is an independent positive regulator of the mechanics and the magnitude of mechanotransduction in cardiomyocytes and skeletal muscle fibers. Given the depth and breadth of our results in striated muscle, and their qualitative alignment with results in other cell types, we found a recent report of an inverse relationship between acetylMT abundance and cardiomyocyte mechanics^15^ rather surprising. However, we believe their use of pharmacologic manipulation on the background of cardiomyopathy, and indirect measures of membrane mechanics, likely contributed to these differences we report here. Furthermore, the strength of our findings are bolstered by measures of passive nanoindentation, active sarcomere mechanics, as well as mechanotransduction dependent ROS signaling; three outcome measures formerly shown effective by our group and others ^8,20^. A further strength is the quantitative similarity of these our measures across mechanistically distinct means of increasing α-tubulin acetylation (i.e, HDAC inhibition vs αTAT1 over-expression). Finally, our results were highly specific to α-tubulin acetylation and occurred independent of alterations in MT density or levels of deTyrMT in both cardiac and skeletal striated muscle.

Our current findings on acetylated α-tubulin, and our works and others on detyrosinated α-tubulin, propose each PTM an independent regulator of striated muscle cell mechanics and mechanotransduction. This independent action would necessitate different mechanisms which appear to be the case. While the detyrosinated α-tubulin c-terminal tail linking to desmin in the sarcomere is the proposed mechanism of how deTyrMTs regulate mechanics and mechanotransduction, acetylated α-tubulin is proposed to promote stability by weakening lateral contacts of microtubule protofilaments^17^. Furthermore, as both PTM’s are elevated in diseased striated muscle, we hypothesize that each act independently towards the pathological increase mechanics and mechanotransduction. Given that a targeted reduction of deTyrMT’s effectively reduces cellular and tissue pathology^8,13,20,25^, we posit that targeting HDAC6 or αTAT may expand our opportunity towards normalizing these pathological changes in diseased striated muscle.

## Material and Methods

### Animal Models and Cell Preparation

Adult C57/BL6 Mice (12-24 weeks) were used to isolate cardiomyocytes and Flexor Digitorum Brevis (FDB) fibers. Cardiomyocytes were isolated using Langendorff retrograde perfusion using 1mg/mL collagenase II^8^, and FDBs were digested overnight in 1mg/mL collagenase A followed by gentle trituration to obtain single cells^8^. Cells were treated with 10µM Tubacin or vehicle (Control) for 2 hours. In a second cohort of animals, FDBs were electroporated with pcDNA3.1 overexpression vector containing GFP (used as control) or αTAT-GFP 4 days prior to enzymatic digestion^8^.

### Western Blots and Immunofluorescence Staining

Isolated cells lysates were run on a polyacrylamide SDS-gel. Protein was transferred to a nitrocellulose membrane and blocked (5% milk). The membrane was probed with primary antibodies for α-Tubulin (ThermoFisher 322588, clone B-5-1-2), acetylated (Sigma T7451, clone 6-11B-1), or detyrosinated tubulin (Abcam ab48389). For immunofluorescence, FDBs or cardiomyocytes were fixed in 4% paraformaldehyde for 20 minutes followed by permeabilization in 0.1% Triton X-100 for 15 minutes. Cells were blocked in SuperBlock PBS (ThermoFisher 37515) for 2 hours. Primary antibodies were used 1:200 overnight at 4°C. Secondary antibodies were incubated at 1:200 for 2 hours. Cells were imaged with Nikon A1R inverted confocal microscope.

### Nano-indentation

FDBs or cardiomyocytes were allowed to settle on a glass bottom dish coated with ECM (Sigma E6909). Cells were indented 1µm with a nano-indenter (Chiaro, Optics11, Netherlands) using a round probe (3 µm radius, 0.044 N/m stiffness). Indentation speeds from 0.5µm/s to 25µm/s were used to calculate the Young’s modulus using the Hertzian contact model. The slowest indentation represents the elastic modulus, while the viscous modulus (ΔE) can be derived from the fastest and slowest indentation^20^.

### Unloaded shortening contractile kinetics

In custom made chambers, FDBs or cardiomyocytes were paced at 1Hz using electric field stimulation (FDB: 600mA 0.2ms, cardiomyocytes: 600mA 2ms). The sarcomere length was recorded using single cell contractile system (Aurora Scientific, Ontario, Canada) and amplitude/kinetics were calculated using a customized MATLAB script.

### Myocyte Attachment and ROS measurements

Isolated FDB fibers were attached to microfabricated glass rod holders (IonOptix), coated with MyoTak as previously described^8^. Mechano-activated ROS production was assayed using Carboxy-H_2_DFFDA (5-(and-6)-carboxy-2′,7′-dihydrofluorescein diacetate; FDBs: 10□μM, 40□min, Cardiomyocytes 6 μM 30 min; ThermoFisher) loaded cells with a 2Hz sinusoidal waveform (10% of resting SL) imposed for 10□s for FDBs, or 2Hz electrical stimulation for cardiomyocytes. Mechanical ROS production was derived from increase in fluorescence during stretch/contractions normalized to basal increase in DFF fluorescence.

### Statistics and Data Availability

Two group comparison was carried out using t-test or Mann-Whitney U-test for parametric and nonparametric data sets respectively. The data is presented as Mean ± SD. The only exception is Stiffness-Indentation velocity relationship curves (Figure 1E and G), where the data comparison was carried out using Two-way analysis of variance (ANOVA) and data is presented as Mean ± 95% CI. The data discussed here, and additional experimental control datasets are available upon request.

## Acknowledgments

This research was supported by NIH T32-AR007592-23 (to AKC); by American Heart Association 19POST34450156 (to HCJ); By NIH U01-HL116321 (to WJL); R01-HL142290 (to WJL and CWW); By NIH R01-AR071618 and R01-AR071614 (to CWW).

